# Coevolution of virulence and immunosuppression in multiple infections

**DOI:** 10.1101/149211

**Authors:** Tsukushi Kamiya, Nicole Mideo, Samuel Alizon

## Abstract

This preprint has been reviewed and recommended by Peer Community In Evolutionary Biology (http://dx.doi.org/10.24072/pci.evolbiol.100043). Many components of host-parasite interactions have been shown to affect the way virulence (i.e., parasite-induced harm to the host) evolves. However, coevolution of multiple parasite traits is often neglected. We explore how an immunosuppressive mechanism of parasites affects and coevolves with virulence through multiple infections. Applying the adaptive dynamics framework to epidemiological models with coinfection, we show that immunosuppression is a double-edged-sword for the evolution of virulence. On one hand, it amplifies the adaptive benefit of virulence by increasing the abundance of coinfections through epidemiological feedbacks. On the other hand, immunosuppression hinders host recovery, prolonging the duration of infection and elevating the cost of killing the host. The balance between the cost and benefit of immunosuppression varies across different background mortality rates of hosts. In addition, we find that immunosuppression evolution is influenced considerably by the precise trade-off shape determining the effect of immunosuppression on host recovery and susceptibility to further infection. These results demonstrate that the evolution of virulence is shaped by immunosuppression while highlighting that the evolution of immune evasion mechanisms deserves further research attention.

## Introduction

The fundamental question of virulence evolution, ‘Why do some parasite strains harm their hosts more than others?’, has been a central focus of evolutionary epidemiology for both its conceptual and applied significance (Ewald, 1994, Read, 1994, Schmid-Hempel, 2011, Méthot, 2012, Alizon and Michalakis, 2015). The adaptive explanation of virulence is typically centred around trade-offs involving virulence and other parasite fitness components, such as transmission and competitiveness in multiple infections (Anderson and May, 1982, Ewald, 1983, van Baalen and Sabelis, 1995, Alizon et al., 2009, 2013). While these trade-off theories explain the evolution of finite non-zero optimal virulence, exactly how much virulence a parasite should evolve depends on a variety of processes (Cressler et al., 2016). For example, host traits (e.g. host immune responses) and their interactions with coevolving parasite adaptations (e.g. parasite immune evasion strategies; Frank and Schmid-Hempel, 2008, Alizon, 2008b, Cressler et al., 2016) are likely to influence the trade-offs. The present theoretical study explores how a parasite immunosuppression strategy, namely the ability of parasites to hinder host recovery, coevolves with virulence.

The ability of parasites to suppress host immunity is ubiquitous in nature (Schmid-Hempel, 2009) and frequently helps maintain chronic infections (Virgin et al., 2009). In humans, for instance, infections by human papillomaviruses (HPVs) and human immunodeficiency virus (HIV) offer two contrasting immune suppression strategies: the former interferes with the cellular machinery to reduce the presentation of viral antigens or impede the interferon response (Doorbar et al., 2012), while the latter infects and lyses T lymphocytes (Levy, 1998). In plant parasites, as well, a variety of mechanisms exist to suppress host defensive responses (Burgyán and Havelda, 2011, Sarmento et al., 2011). Regardless of the specific host-parasite interaction, or mechanism involved, the adaptive benefit for the parasite is realised through prolonged infection duration (Schmid-Hempel, 2009). For the scope of our study, we generalise any parasite adaptation against host immunity that results in lowered host recovery rate as immunosuppression.

In the absence of constraints, it is in the parasite’s best interest to evolve maximal immunosuppression, when immunity serves only to kill parasites. However, lowered host immunity is likely to impose at least one cost to the parasite: an immunocompromised host may be more vulnerable to further infection by conspecific and heterospecific parasites. A meta-analysis by Graham (2008) shows that lowered immune responses, due to the presence of an immunosuppressive helminth, increase microparasite population density within hosts. Furthermore, experimental evidence suggests that immunosuppression could lead to increased host mortality through additional infections by opportunistic parasites (Cornet and Sorci, 2010). Therefore, multiple infections—which are so prevalent that they could be argued to be the rule rather than the exception (Petney and Andrews, 1998, Cox, 2001, Read and Taylor, 2001, Juliano et al., 2010, Balmer and Tanner, 2011) — are likely a key driver of the coevolution between virulence and immunosuppression.

If immunosuppression leads to more multiple infections, one might predict that this should lead to increased virulence. Many theoretical, and some empirical, studies support the notion that within-host competition leads to the evolution of higher virulence (reviewed in Mideo, 2009). Therefore, at the epidemiological level, as the density of coinfected hosts increases, so does the optimal level of virulence (van Baalen and Sabelis, 1995, Choisy and de Roode, 2010). However, given that the benefit of immunosuppression is assumed to be a longer duration of infection, increasing virulence would counteract this effect. Therefore, without a formal model, intuition fails to predict the direction in which virulence evolves when immunosuppression is considered.

To elucidate the coevolutionary dynamics of virulence and immunosuppression, we develop mathematical epidemiology models, in which we assume that the two parasite traits are carried by the same parasite species (as in in van Baalen and Sabelis, 1995). Furthermore, we also investigate how the co-evolved optimal strategy is affected by host background mortality and trade-off concavity determining the effect of immunosuppression on host recovery and susceptibility to further infection.

## The model

We use an evolutionary epidemiology approach based on adaptive dynamics theory (Geritz et al., 1998, Dieckmann et al., 2002, Otto and Day, 2007). We first present the epidemiological model itself, then the evolutionary trade-offs that constrain evolution and finally we show how the (co-)evolutionary analyses are conducted.

### Epidemiological dynamics

We employ a coinfection framework, which allows for coexistence of two parasite strains within a host. Existing coinfection models track either two different resident strains belonging to different species (Choisy and de Roode, 2010) (Fig. 1a), or more simply, a single resident species (van Baalen and Sabelis, 1995) (Fig. 1b). While the two models differ in biological motivations, conceptually, the latter is a special case of the former: the two models are identical if the within-host interactions are the same between the two species (Alizon et al., 2013). Here, we employ the single species model (Fig. 1b) which allows us to study the coevolution of virulence and immunosuppression without making assumptions about how two parasite species are different; thereby requiring fewer parameters. In this model, hosts are divided into three classes: susceptible, singly infected and doubly infected, occurring at densities *S*, *I* and *D* respectively. Following the notation of Table 1, we derive the following system of ordinary differential equations (ODEs) to describe the changes of the resident system over continuous time:

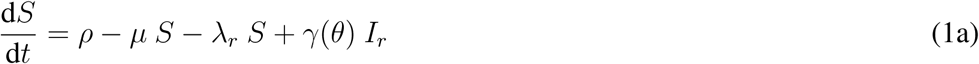

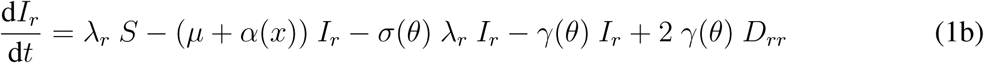

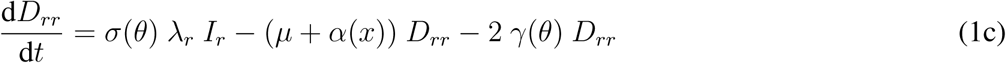

where the subscript *r* denotes the resident parasite strain. In this formulation, there is a constant input of susceptible hosts into the population at the rate *ρ*. Susceptible hosts exit the system through background mortality at the rate *μ*, while infected hosts, both singly and doubly infected individuals, experience additional mortality caused by parasites (i.e., virulence *α*). Susceptible and singly infected hosts acquire infection according to the force of infection λ*_r_* = *βI_r_* + *βD_rr_*, where *β* corresponds to the parasite transmission rate. The host class for double infection by the same strain, *D_rr_* is included in the system for a technical motivation: it is necessary for an unbiased invasion analysis because the mutant strain would gain a frequency-dependent advantage in its absence (discussed in Alizon, 2008a, Lipsitch et al., 2009). We assume that the rate of recovery, *γ*(*θ*), and susceptibility to coinfection, *σ*(*θ*), are functions of immunosuppression, *θ*. Within the existing epidemiological framework, the effect of host immunity can be implicitly accounted for as the rate of recovery (equivalent to the rate of parasite clearance). We assume that hosts recover from infection at a rate *γ*(*θ*), in a stepwise fashion, i.e., doubly infected hosts (*D*) only lose one infection at a time). The key feature of our model is that we assume that singly infected hosts (*I*) suffer an increased risk of contracting a further infection at a rate proportional to a coefficient *σ*(*θ*). We treat the host class *D_rr_* similarly to singly infected hosts *I_r_*, except for the fact that the doubly infected hosts cannot be infected any further.

**Fig. 1:**
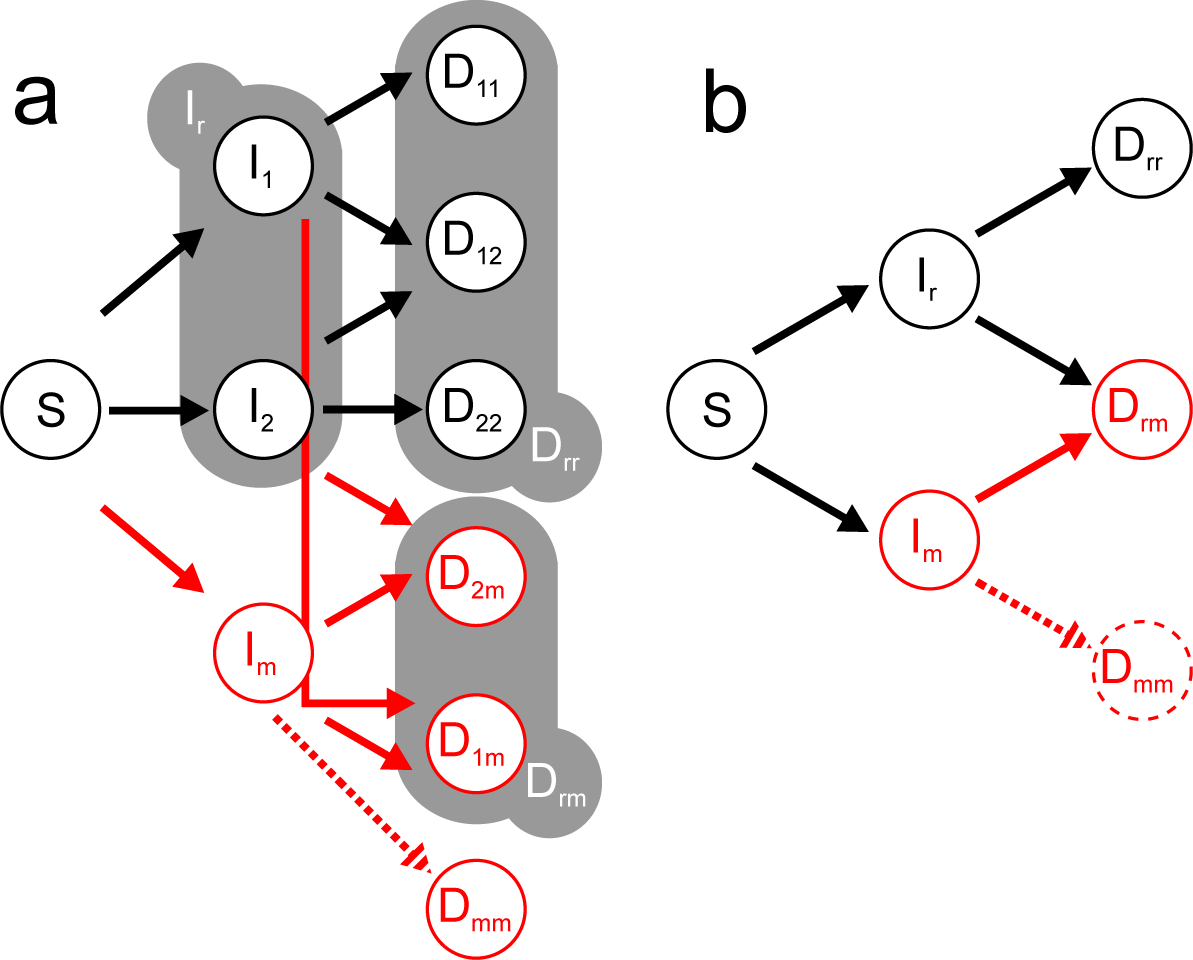
Evolutionary epidemiology model for (a) coinfections by parasites from different species and (b) from same species. In black is the resident system (two strains, one for each species, in (a; labeled 1 and 2) and one strain in (b; labeled r)) and in red are the host classes related to the rare mutant (labeled m). The one species model (b) is a special case of the two species model (a) because the grey bubbles in (a) can be simplified to formulate the one species model (b) when within-host parameters are identical between the two parasite species.

**Table 1:**
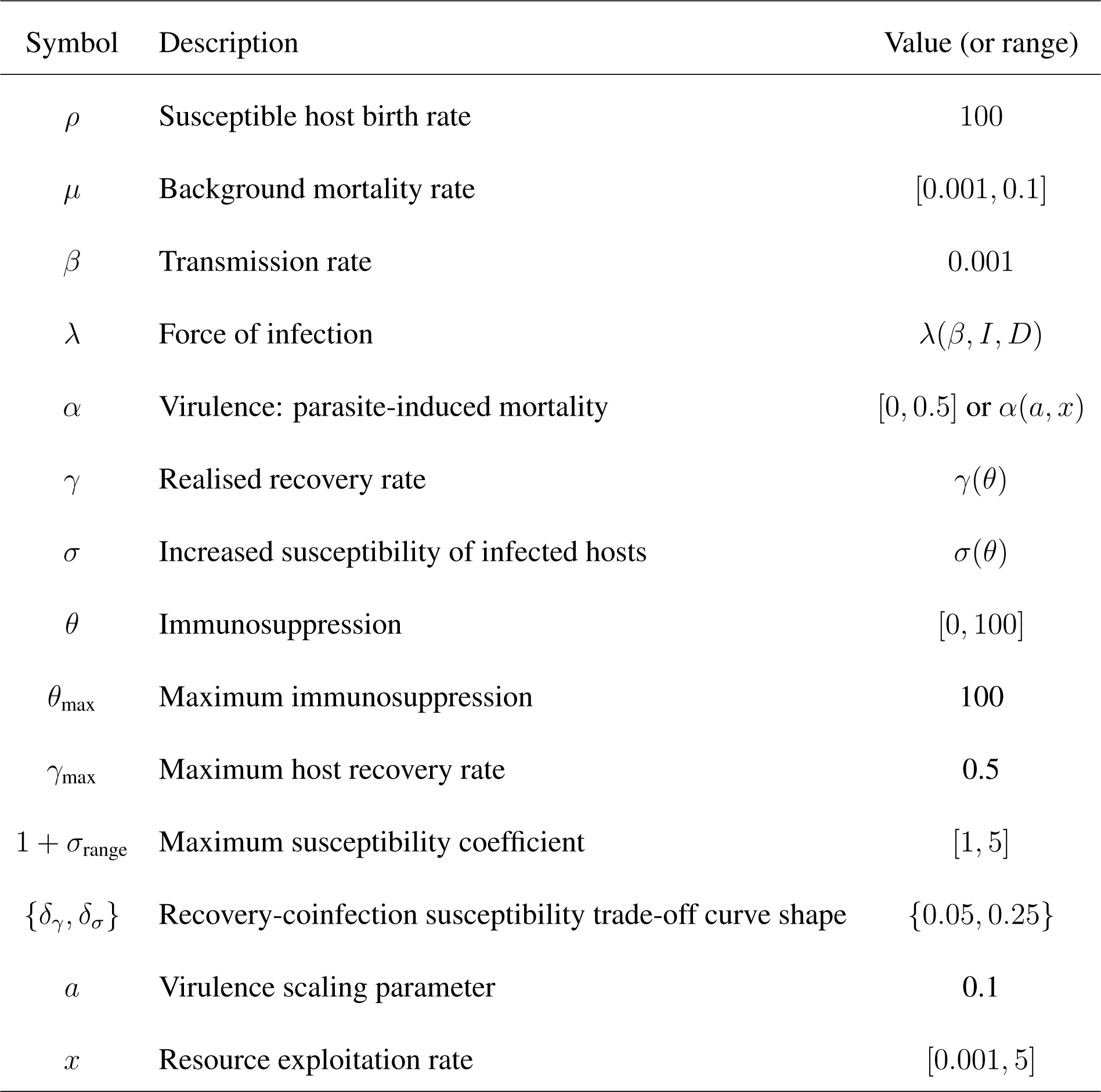
**Parameter notation, description and default values.** Parameter values chosen to sustain non-zero and non-complex equilibria for the resident system and relevant evolutionarily singular strategies. Parameters that are functions of others are indicated with the dependent parameters (or variables) inside parentheses. When we allow only immunosuppression to evolve, virulence, *α*, is a constant; otherwise, *α* evolves as a function of *a* and *x*. Rates are in units of per day.

| Symbol | Description | Value (or range) |
| --- | --- | --- |
| $\rho$ | Susceptible host birth rate | 100 |
| $\mu$ | Background mortality rate | [0.001, 0.1] |
| $\beta$ | Transmission rate | 0.001 |
| $\lambda$ | Force of infection | $\lambda(\beta, I, D)$ |
| $\alpha$ | Virulence: parasite-induced mortality | [0, 0.5] or $\alpha(a, x)$ |
| $\gamma$ | Realised recovery rate | $\gamma(\theta)$ |
| $\sigma$ | Increased susceptibility of infected hosts | $\sigma(\theta)$ |
| $\theta$ | Immunosuppression | [0, 100] |
| $\theta_{\max}$ | Maximum immunosuppression | 100 |
| $\gamma_{\max}$ | Maximum host recovery rate | 0.5 |
| $1 + \sigma_{\text{range}}$ | Maximum susceptibility coefficient | [1, 5] |
| $\{\delta_\gamma, \delta_\sigma\}$ | Recovery-coinfection susceptibility trade-off curve shape | {0.05, 0.25} |
| $a$ | Virulence scaling parameter | 0.1 |
| $x$ | Resource exploitation rate | [0.001, 5] |

### Within-host processes and resulting trade-offs

It is commonly assumed that virulence (i.e., parasite-induced host mortality) correlates with the extent of parasite resource exploitation. Adaptive benefits of resource exploitation include the positive correlation with transmission (Fraser et al., 2007, de Roode et al., 2008, Råberg, 2012), and a within-host competitive advantage in coinfection (de Roode et al., 2005, Bell et al., 2006, Ben-Ami et al., 2008, Zwart et al., 2009). Here, we focus on the latter adaptive benefit to study the evolution of virulence and immunosuppression. We assume that virulence (*α*) increases linearly with the level of resource exploitation by a parasite (*x*), such that *α*(*x*) = *a x*, where *a* is a proportionality constant (we explore a transmission-virulence trade-off in the Supplementary Information 2). We then assume that finding themselves in a doubly infected host is inherently costly for parasites due to exploitation competition between coinfecting strains (Mideo, 2009, Schmid-Hempel, 2011), and that more virulent strains are more competitive in multiple infections:

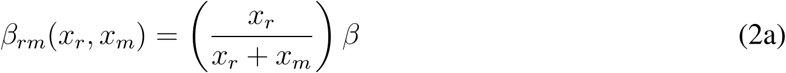

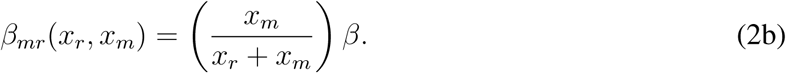

There is ample empirical evidence that immunosuppression benefits the parasites by prolonging infections (reviewed in Schmid-Hempel, 2008), and lowered host immunity would increase the susceptibility to multiple infections (Palefsky and Holly, 2003, Rockstroh and Spengler, 2004, Cornet and Sorci, 2010). Thus, the key trade-off in our model is between infection duration and susceptibility to coinfections (both being mediated by immunosuppression). We, therefore, assume a trade-off between the rate of recovery, *γ*(*θ*), and additional susceptibility of infected hosts to coinfection, *σ*(*θ*), by making them both functions of immunosuppression intensity, *θ*. It is conceivable for the decline of recovery rate and the increase of additional susceptibility to either accelerate or decelerate with increasing immunosuppression. Because the trade-off shape typically matters for evolutionary dynamics (Bowers et al., 2005, Kisdi, 2006) and little is known from empirical data, we explore the trade-offs involving recovery and susceptibility as both accelerating and decelerating functions of immunosuppression. The parameters *δ_γ_* and *δ_σ_* control the degree of concavity of the effect of immunosuppression on recovery and increased susceptibility, respectively (eq. 3; Fig. S1).

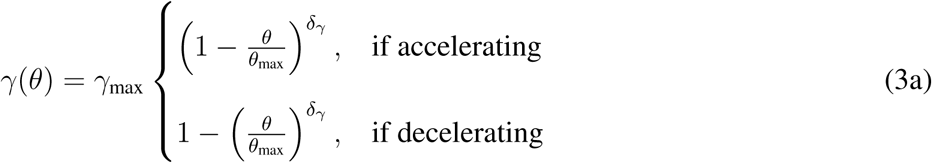

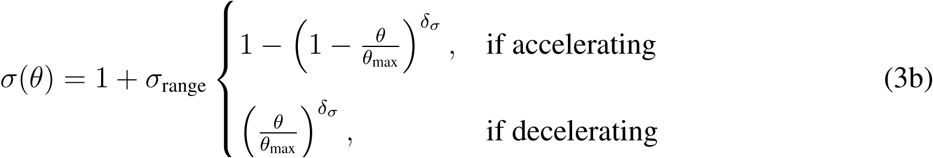

With these functions, we assume that the realised recovery rate, *γ*(*θ*), decreases as a function of immunosuppression such that it equals the intensity of host immunity, *γ*_max_, in the absence of immunosuppression and approaches 0 as immunosuppression approaches *θ*_max_. We also assume that the proportional gain in susceptibility to a further infection, *σ*(*θ*), elevates the force of infection experienced by an immunosuppressed singly infected host by up to 1 + *σ*_range_ fold at the upper limit of immunosuppression (when *θ* = *θ*_max_). Because it is commonly assumed that the pay-off of a beneficial trait saturates, we set the recovery trade-off as decelerating at default. We set the default susceptibility trade-off as accelerating to further emphasise the difference between beneficial and costly traits.

### Evolutionary analyses

#### The mutant systems

We carry out an invasion analysis investigating perturbation of the resident state by adding a rare mutant strain, the densities and traits of which are denoted with subscript *m* (Fig. 1b). For the evolution of immunosuppression, the dynamics of the mutant strain are summarised in the following system of ODEs:

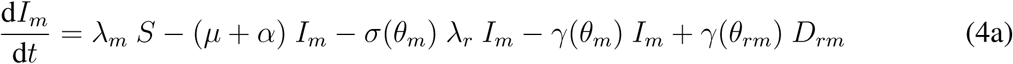

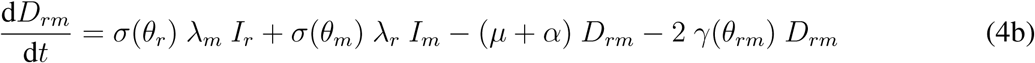

where *λ_r_* = *βI_r_* + *βD_rr_* + *β_rm_D_rm_* and *λ_m_* = *βI_m_* + *β_mr_D_rm_*. For simplicity we assume that the order of infection does not matter so that *D_rm_* is identical to *D_mr_*. We neglect hosts infected twice by the mutant strain (which would be *D_mm_*) because it is unlikely that the same host gets infected twice by a rare mutant. Recovery from *D_rm_* can be achieved through either clearing a resident or a mutant parasite. Other aspects of demographic changes of the mutant system are identical to the resident system described above.

We assume that the level of immunosuppression in coinfection is the average between the resident and mutant strain, i.e.,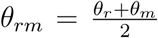. For virulence evolution, we assume that the only within-host interaction between coinfecting parasites is competition for the shared host resources. Therefore, we also calculate the overall virulence of coinfection as the average of the two strains, i.e.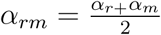.

The mutant dynamics for virulence evolution are governed by

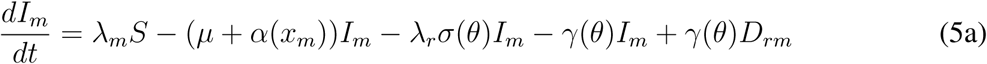

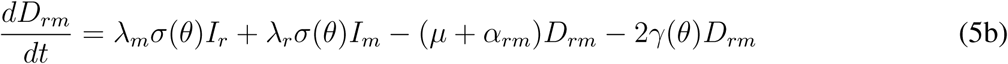

where *λ_r_* and *λ_m_* are the force of infection for the resident and mutant, respectively, defined here as *βI_r_* + *βD_rr_* + *β_rm_D_rm_* and *βI_m_* + *β_mr_D_rm_*. We again assume the trade-offs between recovery and coinfection susceptibility as functions of immunosuppression in this model.

#### Adaptive dynamics

The fate of a rare mutant strain is determined by its fitness function (here denoted *R_θ_m__* and *R_α_m__*, respectively), that is, the ability to spread through a host population already infected with a resident parasite (Geritz et al., 1998, Dieckmann et al., 2002). In the continuous time scale, the mutant parasite invades and replaces the resident if the mutant fitness, calculated as the dominant eigenvalue of the Jacobian matrix of the mutant system, is positive (Otto and Day, 2007). The expressions for the invasion fitness of a rare mutant — with respect to immunosuppression and virulence (*R_θ_m__* and *R_α_m__*, respectively) — emerging in a population infected by a resident strain are:

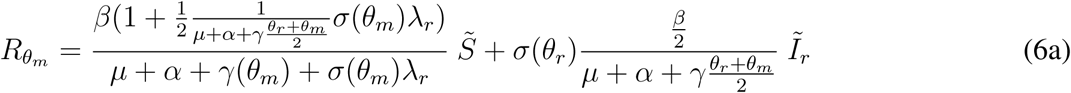

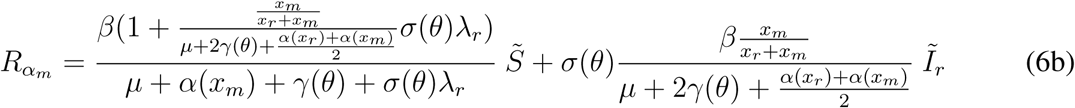

Consequently, an evolutionarily singular strategy can be found where the change of the invasion fitness ceases with respect to the evolving trait. For example, an evolutionarily singular strategy of immunosuppression (denoted *θ*^*^) can be found when *θ*^*^ is an extremum of *R_θ_m__*:

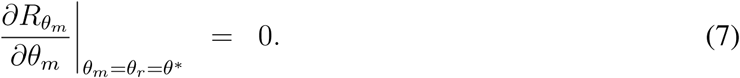

The properties of a singular strategy can then be assessed by the second derivatives of *R_θ_m__*. Following the notations used by Geritz et al. (1998), here we denote the second derivatives of *R_θ_m__* with respect to the resident and mutant strain with *a* and *b*:

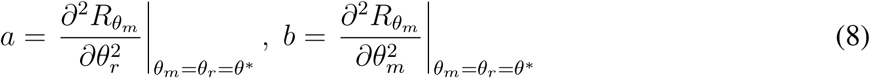

The convergence stable ES (i.e. the strategy towards which selection drives the population and that is also non-invasible by mutants; i.e., evolutionarily stable and convergence stable, or the continuously stable strategy, CSS *sensu* Eshel (1983)) condition is satisfied when *b* < 0 and *a* − *b* > 0. The first condition states that the mutant fitness is at a local maximum and hence evolutionarily stable and the second condition implies no mutant invasion is possible at the point, meaning convergence stable (Geritz et al., 1998). Various other possible configurations of evolutionarily and convergence stability are discussed in Geritz et al. (1998).

#### Coevolution of virulence and immunosuppression

We graphically identified the coevolutionarily singular state as the intersection between the singular state of immunosuppression and virulence (Choisy and de Roode, 2010, Alizon, 2013). When this intersection is both convergence and evolutionarily stable, it can be interpreted as the coevolutionarily stable strategy (Maynard Smith, 1982, Marrow et al., 1996, Dieckmann et al., 2002). The conditions for coevolutionary stability are given in detail by (Abrams et al., 1993, Marrow et al., 1996). In brief, the stability of each co-evolving trait is neither sufficient nor necessary, and there is no simple set of criteria that guarantees local asymptotic stability. We explore the coevolution of the two traits across different extrinsic mortality conditions and immunosuppression trade-off concavity.

## Results

### Virulence evolution

We first assume that the level of immunosuppression is constant and infer the virulence level towards which the parasite population evolves, that is the evolutionarily stable virulence (ESV). We find that the higher the immunosuppression, the higher the ESV (grey curve in Fig. 2a). Because immunosuppression renders infected hosts more susceptible to further infections, it consequently increases the relative abundance of doubly infected hosts (Fig. 2e). This favours more virulent parasites due to within-host competition (see equation 2).

**Fig. 2:**
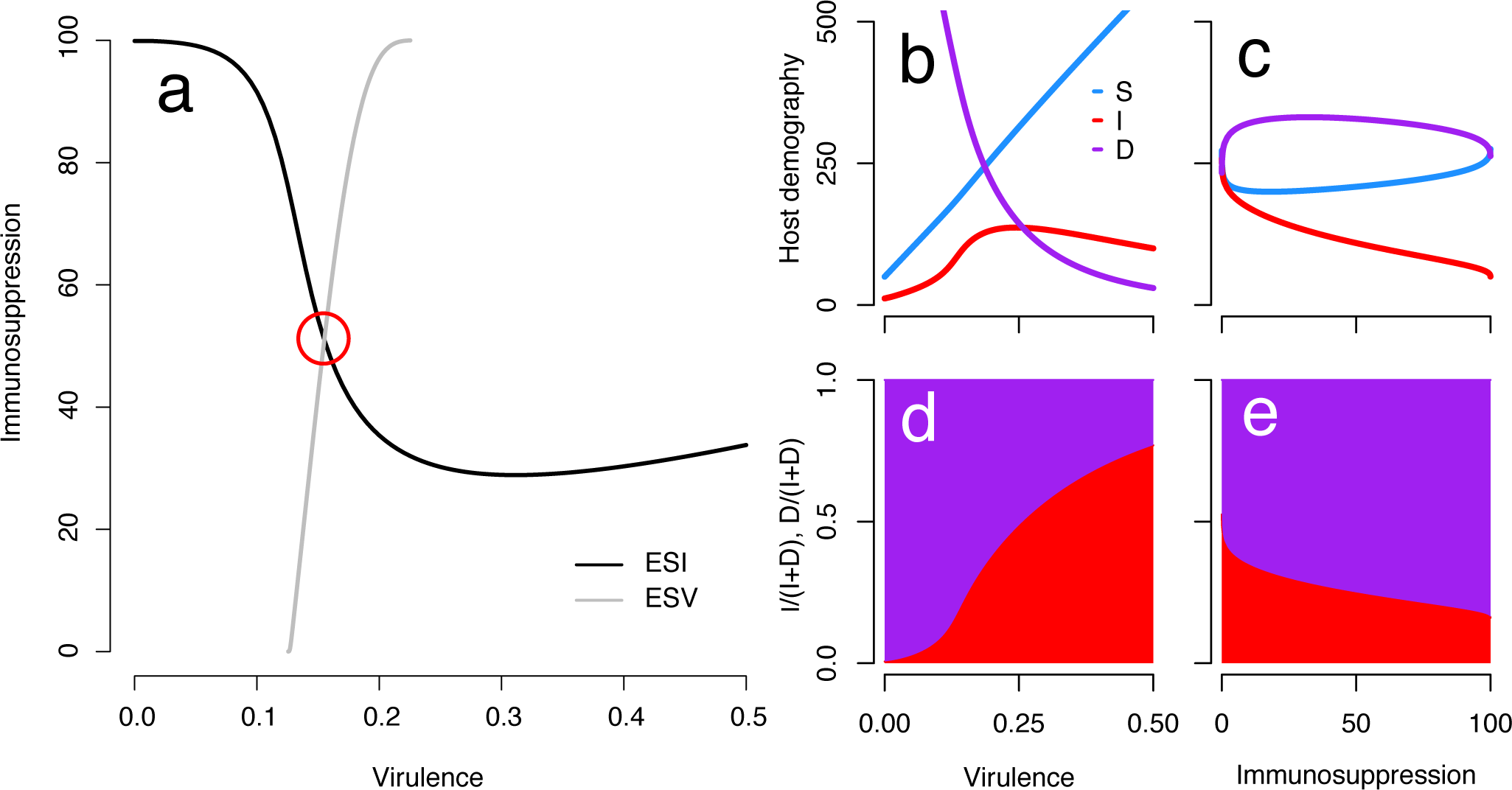
(a) Evolutionarily stable immunosuppression (ESI; black) and virulence (ESV; grey) against fixed values of the other trait. The co-evolutionarily stable strategy (co-ESS) of the two traits occurs at the intersection of the two lines, indicated by the red circle. The immunosuppression trade-offs for the recovery rate and additional susceptibility were decelerating and accelerating, respectively with shape parameters *δ*_*γ*_ = 0.05 and *δ*_*σ*_ = 0.25. The equilibrium population size of the three host classes — susceptible (*S*; blue), singly infected (*I*; red) and doubly infected (*D*; purple) — underlying the ESI over a range of of virulence and the ESV over a range of immunosuppression values is presented in (b) and (c). The relative abundances of singly (red) and doubly (purple) infected host are plotted in (d) and (e).

### Immunosuppression evolution

We then set the virulence to a constant value and study whether parasite immunosuppression evolves towards an evolutionarily stable strategy (i.e., evolutionarily stable immunosuppression, or ESI; black curve in Fig. 2a). We find that ESI decreases with virulence at first, but it increases again when virulence is high enough. The initial decrease can be attributed to two non-mutually exclusive processes. First, the benefits gained by increasing immunosuppression (i.e., slower host recovery) are reduced as virulence increases since the duration of infection decreases. Note that ESI similarly decreases as host mortality increases (Fig. 3a). Second, the decreasing pattern may originate from demographic feedbacks: increasing virulence reduces coinfections, therefore parasites reap the benefit of immunosuppression in reduced recovery without paying the cost of contracting further infections. Therefore, the initial decrease in ESI with virulence is also likely mediated by the falling fraction of multiple infections (Fig. 2d).

**Fig. 3:**
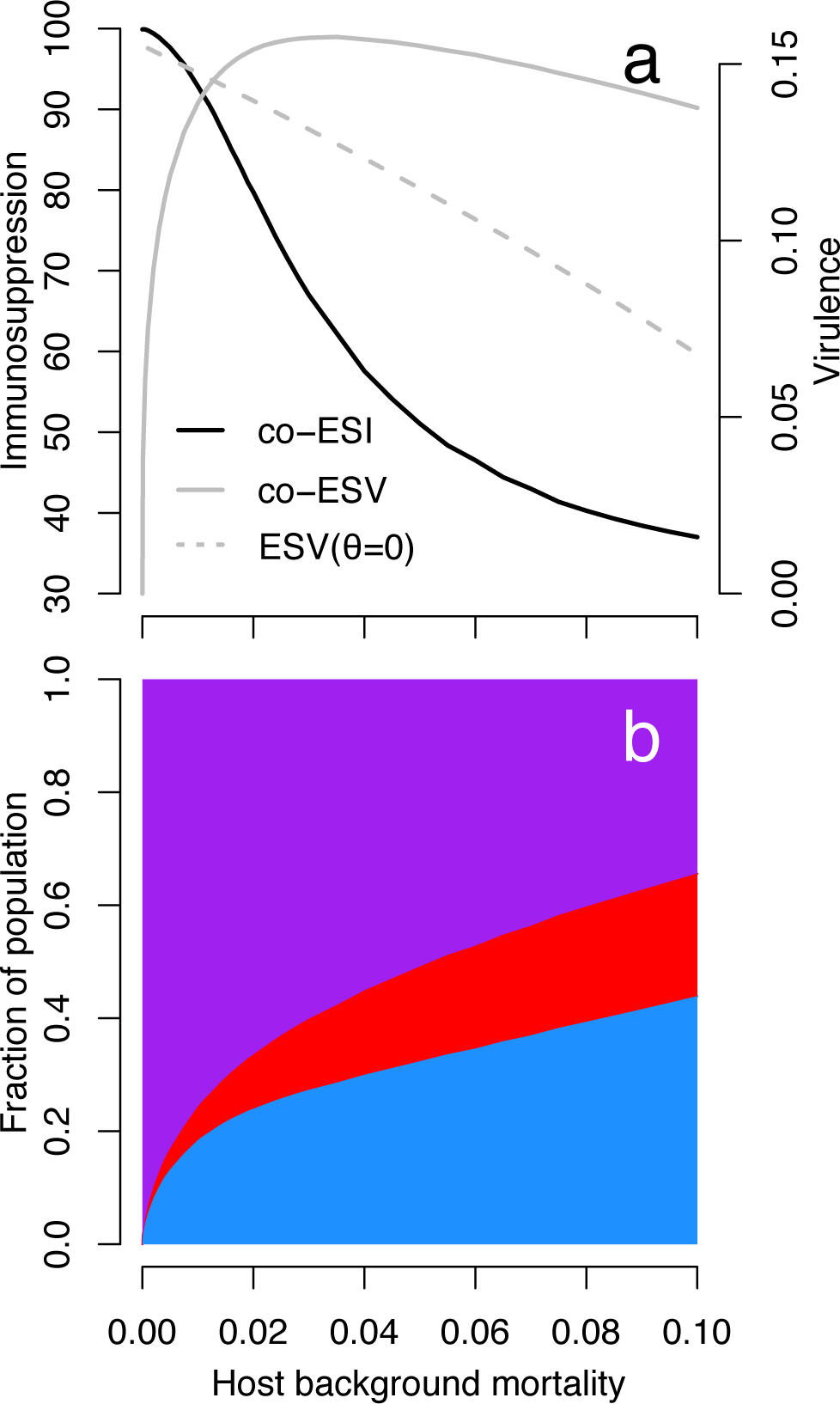
(a) Coevolutionarily stable immunosuppression (co-ESI; black) and virulence (co-ESV; solid grey) strategies, and evolutionarily stable virulence strategy in the absence of immunosuppression (ESV (*θ* = 0); dashed grey) against host background mortality and (b) the fraction of population of the three host classes — susceptible (*S*; blue), singly infected (*I*; red) and doubly infected (*D*; purple) at the coevolutionarily stable state.

We also find that the ESI increases with virulence when virulence is high enough. As the lifespan of an infected host decreases due to high parasite-induced morality, it becomes unlikely for a host to survive a single infection long enough to get infected again. At this point, coinfections are sufficiently rare (Fig. 2d) that highly immunosuppressive parasites would rarely suffer the cost of immunosuppression in contracting further infections. Taken together, focusing on the prevalence of coinfections alone is not enough to predict how ESI will evolve.

### Coevolution of virulence and immunosuppression

The co-ESS is found at the intersection between the two curves in Figure 2. For our default parameters, this occurs at intermediate values of immunosuppression and virulence. We now investigate how changes in host mortality and the trade-off shapes determining the effect of immunosuppression on host recovery and susceptibility to further infection affect this co-ESS. We first explore how the co-ESS varies with respect to the rate of host background mortality. We find that co-ES immunosuppression (co-ESI) always decreases with host background mortality (black line in Fig. 3a), in accord with the intuition that immunosuppression represents a lost investment if the host dies too rapidly.

In the absence of immunosuppression, as found in previous models (van Baalen and Sabelis, 1995, Gandon et al., 2001), the optimal virulence decreases with host background mortality because the higher the mortality, the lesser the chance of coinfection from which the benefit of virulence is realised (dashed grey line in Fig. 3a purple area in b). In contrast, we find that virulence coevolving with immunosuppression (co-ESV), peaks at an intermediate value of background mortality (solid grey line in Fig. 3a). Considering an extreme case in which the host never dies through background mortality (i.e., *μ* = 0), the best strategy for the parasite is to evolve avirulence and maximise immunosuppression so that the host remains infected forever (Fig. 3a). This scenario can be interpreted as an alignment of interest between resident and mutant strains as the benefit of keeping the host alive longer appears to outweigh the adaptive advantage of being competitively dominant. With zero mortality and maximum immunosuppression, a parasite’s fitness is infinite: any mutant with some virulence will have a finite fitness (because it will kill its host in single and double infections). Intuitively, this avirulent strategy can also invade because in absence of the virulent strain, the fitness is always maximised and the advantage reaped by the virulent one in coinfection is not enough to overcome the cost of killing its singly-infected hosts. This cost of killing the host in single infection relaxes as mortality increases, leading to a steep increase in virulence. The eventual decrease in virulence is consistent with the evolution of virulence in the absence of immunosuppression (dashed grey line in Fig. 3a)(van Baalen and Sabelis, 1995, Gandon et al., 2001).

Little is known about how immunosuppression impacts host recovery and susceptibility to further infection. Therefore, we also explored the sensitivity of our co-ESS results to the qualitative shape of the immunosuppression trade-off and the extent of its concavity using parameters, *δ_σ_* and *δ_γ_*. We find that evolution moves away from the singular strategy when the recovery concavity is highly accelerating (Fig. 4a) meaning that in this case immunosuppression is either maximised or minimised depending on the initial conditions. Furthermore, we find that immunosuppression is maximised for a large area of the near-linear and decelerating recovery trade-off space, *δ*_*γ*_. Intermediate ESI levels are observed for highly decelerating recovery, *δ*_*γ*_. Overall, this suggests that there is a tendency for parasites to specialise in immunosuppressing their host or to completely avoid doing so, and knowledge of the recovery function appears particularly important for predicting immunosuppression evolution. For virulence, the concavity of the susceptibility function (*δ*_*σ*_) has the strongest quantitative effect, with decelerating trade-offs leading generally to higher co-ESV. As in the rest of this model, since the only benefit associated with virulence is increased competitiveness in a coinfected host, the co-ESV is an indicator of the importance of this competition in the parasite’s life cycle.

**Fig. 4:**
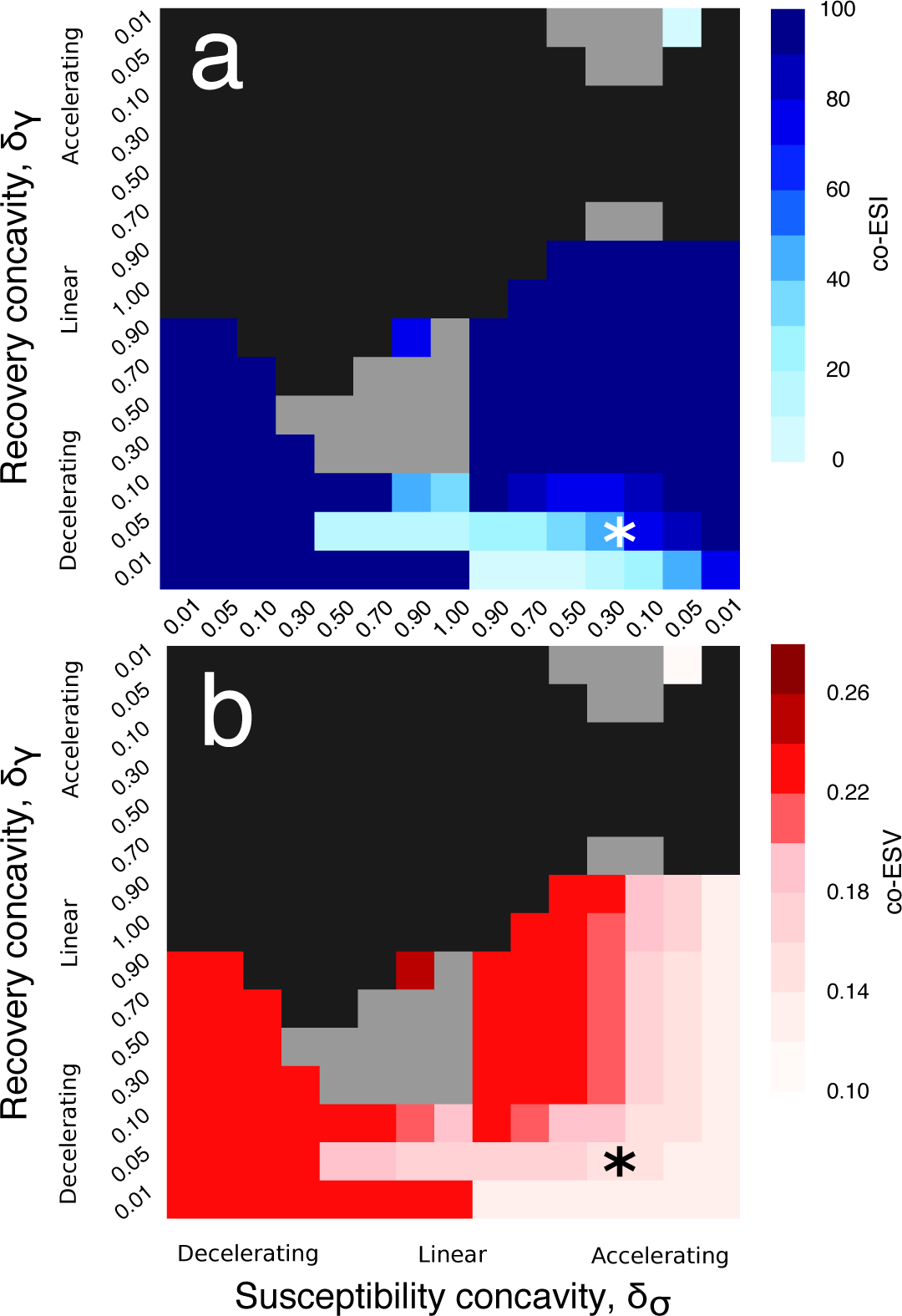
The trade-off concavity influences the coevolutionary outcome. The shade of blue and red indicates the coevolutionarily stable strategy value of (a) immunosuppression and (b) virulence, respectively. The asterisk (*) indicates the default set of trade-off parameters explored in Figure 2 and 3. The dark grey areas indicate that the coevolutionarily singular strategy is an invasive repeller. The light grey squares indicate that the outcome of coevolutionary stability depends on the details of the rate and variance of mutational inputs of the two coevolving traits.

## Discussion

Host immune responses present a major challenge for parasites and, so, establishing a successful infection often depends upon a parasite’s ability to evade host immunity (Schmid-Hempel and Frank, 2007, Kerr et al., 2017). Despite its ubiquity among all major groups of parasitic organisms (Schmid-Hempel, 2009), the effect of immunosuppression on virulence evolution has largely been overlooked (but see Koella and Boete, 2003, Hurford and Day, 2013). We modelled immunosuppression through its joint effect on host recovery and susceptibility to coinfection in an attempt to understand epidemiological forces driving the coevolution of virulence and immunosuppression.

We found that immunosuppression increases the optimal parasite exploitation by creating more coinfections, in which more competitive (and hence more virulent) strains are favoured. On the other hand, the evolution of immunosuppression is driven by the balance between the benefit conferred by immunosuppression to evade clearance from the host and the associated cost of contracting further infections, which introduce a competitor for limited host resources. Because virulence simultaneously decreases both the benefit (by killing hosts faster) and the cost (by reducing the risk of coinfection), its effect on the optimal immunosuppression is nuanced — increasing virulence can both increase or decrease the optimal immunosuppression depending on the baseline virulence of the parasite.

We then investigated the change in coevolutionarily optimal strategies of the two traits over host background mortality. We find that mortality decreases the coevolutionarily stable level of immunosuppression, which is a lost investment when hosts die too fast anyway. In the absence of immunosuppression, we expect the optimal virulence to consistently decrease with host background mortality because, again, investing in competitive ability (with which virulence correlates) is wasted when coinfections are rare (van Baalen and Sabelis, 1995, Gandon et al., 2001). When coevolving with immunosuppression, however, we find that evolutionarily stable virulence peaks for an intermediate level of host mortality. This stems from the fact that for low host mortality, the coevolutionarily optimal parasite strategy is to prolong the duration of infection by simultaneously maximising immunosuppression and minimising virulence.

In light of our theoretical model, we can formulate testable predictions. In *Daphnia*, for example, the rate of host background mortality can be experimentally manipulated and its effect on virulence evolution of microsporidian parasites can be quantified (Ebert and Mangin, 1997). Microsporidians are common eukaryotic parasites of many animals including *Daphnia*, which often harbour multiple infections (Ebert, 2005). In their mosquito host, microsporidians have been suggested to suppress host immunity by manipulating the production pathway of a host immune defence molecule (nitric-oxide, NO), which is part of the innate immune system conserved in all animals (Biron et al., 2005). Conveniently, the production of NO can also be experimentally enhanced and blocked, making it possible to investigate the effects of manipulating host immune intensity (Rivero, 2006). While our model predicts the coevolution of virulence and immunosuppression is likely influenced by the precise shape of the trade-offs determining the cost and benefit of immunosuppression, there is a dearth of empirical data with which to calibrate these curves. The *Daphnia* system may offer an opportunity to characterise immunosuppression trade-offs and advance the understanding of the role of immunosuppression in virulence evolution.

A natural extension to the model of coinfection by the same species (van Baalen and Sabelis, 1995) is the model that accommodates two distinct resident parasite species, each of which can be challenged by a mutant (Choisy and de Roode, 2010). Under the different species model, two co-evolving traits (e.g., immunosuppression and virulence) could be carried by two separate parasite species, which better reflect the reality for some immunosuppressing parasites, e.g., the immunosuppressing capabilities of HIV render the host susceptible to the virulence induced by opportunistic infections. Similarly, in an amphipod system, Cornet and Sorci (2010) show that immunosuppressive parasites elevate host mortality by promoting opportunistic pathogen infections. Furthermore, there is evidence that pathological severity of malaria infection can be amplified through immunosuppression caused by helminths, which are common parasites in malaria prevalent tropical regions (Graham et al., 2005). That being said, considering multiple species would force us to revisit our assumption that more virulent mutants are more competitive than their resident at the within-host level. Indeed, this assumption has recently been shown to hold for a variety of within-host processes, but only if the mutant traits are close to that of the resident (Sofonea et al. in prep). Therefore, adding more details about the within-host interactions, e.g., via a nested model (Mideo et al., 2008), seems necessary to study coinfection by different species.

In the present model, we assumed no direct link between immunosuppression and virulence. However, immune evasion strategies of bacteria and viruses have been empirically linked to a range of pathological effects (Casadevall and Pirofski, 2003, Monack et al., 2004, Stanford et al., 2007). On the other hand, immunosuppression may decrease immunopathology which can, therefore, reduce host mortality, as shown experimentally using rodent malaria infections (Long et al., 2008, Long and Graham, 2011). In fact, helminth therapy, which involves deliberate ingestion of parasitic worms, takes advantage of the parasite’s ability to mediate host immunity and has been successful in countering inflammations caused by immune-mediated diseases (Day et al., 2007, Elliott and Weinstock, 2009, Summers et al., 2003).

The only cost of immunosuppression we assumed is indirect (coinfection facilitation), however, the production of immunosuppressive compounds could impose a direct fitness cost to individual parasites. At the within-host level, immunosuppression would, therefore, be seen as a public good since parasites that do not invest in it can still reap the benefits (Diard et al., 2013, Rundell et al., 2016). In fact, our model predicts that invasive repellers are common while coexistence of two strains with extreme immunosuppression strategies (i.e., zero and maximum immunosuppression) is always possible regardless of trade-off concavity (figure not shown). These findings suggest that it may be common for some strains to specialise in immunosuppressing and others in exploiting these immunosuppressed hosts.

Understanding how host immunity and the corresponding parasite immune evasion strategies affect virulence evolution is a key challenge for contemporary evolutionary epidemiology (Frank and Schmid-Hempel, 2008). Our results demonstrate that immune evasion mechanisms are among the major forces shaping virulence evolution at the between-host level. Future theoretical studies may focus on multi-species epidemiological dynamics, direct trade-offs between immunosuppression and virulence and life-history perspectives.

## Acknowledgements

We thank Sébastien Lion, Stéphane Cornet, Philip Agnew, Matthew Hartfield, Yannis Michalakis and Mircea Sofonea for comments and discussions and Céline Devaux and Katie O’Dwyer for comments on an earlier draft. We are also grateful for valuable comments provided by Sara Magalhaes and two anonymous reviewers through the Peer Community in Evolutionary Biology.

